# Cytolysin-positive *Enterococcus faecalis* is elevated in patients with chronic alcoholic pancreatitis

**DOI:** 10.1101/2021.11.30.468848

**Authors:** Clara Perrin, Vinciane Rebours, Nicolas Trainel, Cosmin Sebastian Voican, Gabriel Perlemuter, Anne-Marie Cassard, Dragos Ciocan

## Abstract

**Introduction:** Patients with alcoholic hepatitis have an increase in cytolysin-producing *Enterococcus faecalis* that correlates with disease severity and mortality.

**Aim:** To determine whether patients with chronic alcoholic pancreatitis have an elevated abundance of cytolysin-producing *E. faecalis*.

**Methods:** Quantification by qPCR of cytolysin-producing *E. faecalis* in controls and patients with alcoholic hepatitis or pancreatitis.

**Results:** Patients with alcoholic pancreatitis had a higher proportion of intestinal cytolysin-positive *E. faecalis* than healthy controls and patients with alcoholic hepatitis.

**Conclusion:** **C**ytolytic *E. faecalis* may also be involved in this other alcohol-related complication and benefit from targeted microbiota editing strategies.

## INTRODUCTION

Chronic, excessive alcohol consumption leads to several diseases, including alcoholic hepatitis (AH) and chronic alcoholic pancreatitis (CAP). These diseases are life-threatening complications of chronic alcohol consumption, with a high rate of mortality, and they have no specific treatment, except alcohol withdrawal. Several studies have shown that the intestinal microbiota plays a role in the pathogenesis of these two diseases and that patients with CAP or AH display altered and specific gut dysbiosis (1,2). Among the differences observed in the intestinal microbiota, we previously reported a higher relative abundance of *Enterococcus* in patients with CAP (2).

A recent study by Duan et al. found that patients with AH have an elevated relative abundance of a particular type of fecal *Enterococcus faecalis* that produces a bacteriocin, cytolysin, which causes hepatocyte death. This bacteria is associated with more severe clinical outcomes and increase mortality in these patients (3). However, it was not clear in this study whether the alcoholic patients with alcoholic hepatitis that were included had any other alcohol-related complications. This could be of major interest, as the microbiome-editing strategy proposed by the authors could also be extended to other alcohol-induced complications if the relative abundance of this strain of *Enterococcus faecalis* is also elevated in such complications. We thus aimed to determine whether patients with chronic alcoholic pancreatitis also exhibit an elevated relative abundance of cytolysin-producing *Enterococcus faecalis* and compare their profile to that of patients with alcoholic hepatitis.

## METHODS

Three groups of patients were included in the study: healthy controls (HC) (n = 28), patients with AH and without CAP or acute pancreatitis (n = 27) and patients with CAP and without AH (n = 24). Fecal samples were frozen at -80°C and bacterial DNA was extracted as previously described (4). We amplified three bacterial genes: total bacterial 16S, *E. faecalis* 16S, and *E. faecalis cytolysin L (cylL*_*L*_) (see Supplementary Material for details).

## RESULTS

The patient characteristics are summarized in Table 1. We included patients with chronic alcohol consumption and two types of alcohol-related complications, AH without CAP and CAP without AH. Patients with CAP had a significantly higher abundance of fecal *E. faecalis* than healthy controls (HC) (p < 0.0001) and AH patients (p = 0.0004) (Figure 1A). There was no difference in the relative abundance of *E. faecalis* between AH patients and HC (Figure 1A). Genomic DNA of *E. faecalis* was detected in 96% (23/24) of CAP patients, 89% (25/28) of HC (p > 0.05), and 85% (23/27) of AH patients (p > 0.05) (Figure 1B). CAP patients were more frequently cytolysin-positive (23/24, 96%) than HC (11/28, 39%, p < 0.001) or AH patients (13/27, 48%, p < 0.001, Figure 1C). In addition, the relative abundance of cytolysin-positive *E. faecalis* was elevated in CAP and AH patients (p = 0.0006 and p= 0.0067) but not HC (Figure 1D). The relative abundance of cytolysin-positive *E. faecalis* did not correlate with pancreatic disease severity, neither in terms of biological (inflammation and albumin levels) nor radiological (necrosis and inflammation) severity criteria.

**Table 1.** Clinical characteristics of the three groups of patients.

|  | Chronic alcoholic pancreatitis (CAP, n = 24) | Severe alcoholic hepatitis (AH, n = 27) | Healthy controls (HC, n = 28) |
| --- | --- | --- | --- |
| **Age (years)\*\*\*** | 51.4 ± 9.8 | 52.5 ± 11.5 | 34.9 ± 11.6 |
| **Sex (male/female)\*\*\*** | 21/3 | 23/4 | 12/16 |
| **BMI (kg/m<sup>2</sup>)\*** | 22.3 ± 3.3 | 26 ± 5.2 | 23 ± 3.6 |
| **Alcohol intake (g/day) \*** | 143.8 ± 90 | 106 ± 65.5 |  |
| **Duration of intake (years)\*\*** | 13 ± 3.9 | 21 ± 10.8 |  |
| **Smoking (%)** | 22 (92) | 17 (65) |  |
| **Type 2 diabetes (%) \*\*** | 10 (42) | 2 (8) |  |
| **PPI use (%)** | 10 (45) | 8 (30) |  |
| **CRP (mg/L)\*** | 37.4 ± 72..1 | 31.88 ± 25..3 |  |
| **AST (U/L)\*\*\*** | 38 ± 34..3 | 242 ± 495..7 |  |
| **ALT (U/L)** | 43.9 ± 43..9 | 66.4 ± 88..4 |  |
| **Total bilirubin (μmol/L)\*\*\*** | 25 ± 68 | 210.5 ± 172 |  |
| **GGT (U/L)\*\*\*** | 182..9 ± 287..9 | 392.8 ± 314..5 |  |
| **Glycaemia (mmol/L)\*\*\*** | 7.3 ± 1.9 | 5.4 ± 1 |  |
| **Serum albumin (mg/dL)** | 31 ± 7.8 | 26.9 ± 4.8 |  |
| **Platelets (x10<sup>9</sup>/L)\*\*\*** | 325 ± 129 | 109 ± 88 |  |
| **Prothrombin time (%)\*\*\*** | 94 ± 13.9 | 39 ± 13.5 |  |
| **Cirrhosis (%)\*\*\*** | 0 | 27 (100) |  |
| **Liver biopsy (%)\*\*\*** | 0 | 27 (100) |  |
| **Maddrey score** |  | 55.9 ± 21.3 |  |
| **MELD score** |  | 21.5 ± 6.5 |  |
| **Exocrine pancreatic insufficiency (%)\*\*** | 6 (25) | 0 |  |

Data is expressed as the mean ± SD for continuous variables and n (%) for discrete variables. PPI: proton pump inhibitors, CRP: C-reactive protein, AST: aspartate transaminase, ALT: alanine transaminase, GGT: gamma-glutamyl transpeptidase, MELD: Model for End-Stage Liver Disease. \*p<0.05, \*\*p<0.01, \*\*\*p<0.001

**Figure 1.**
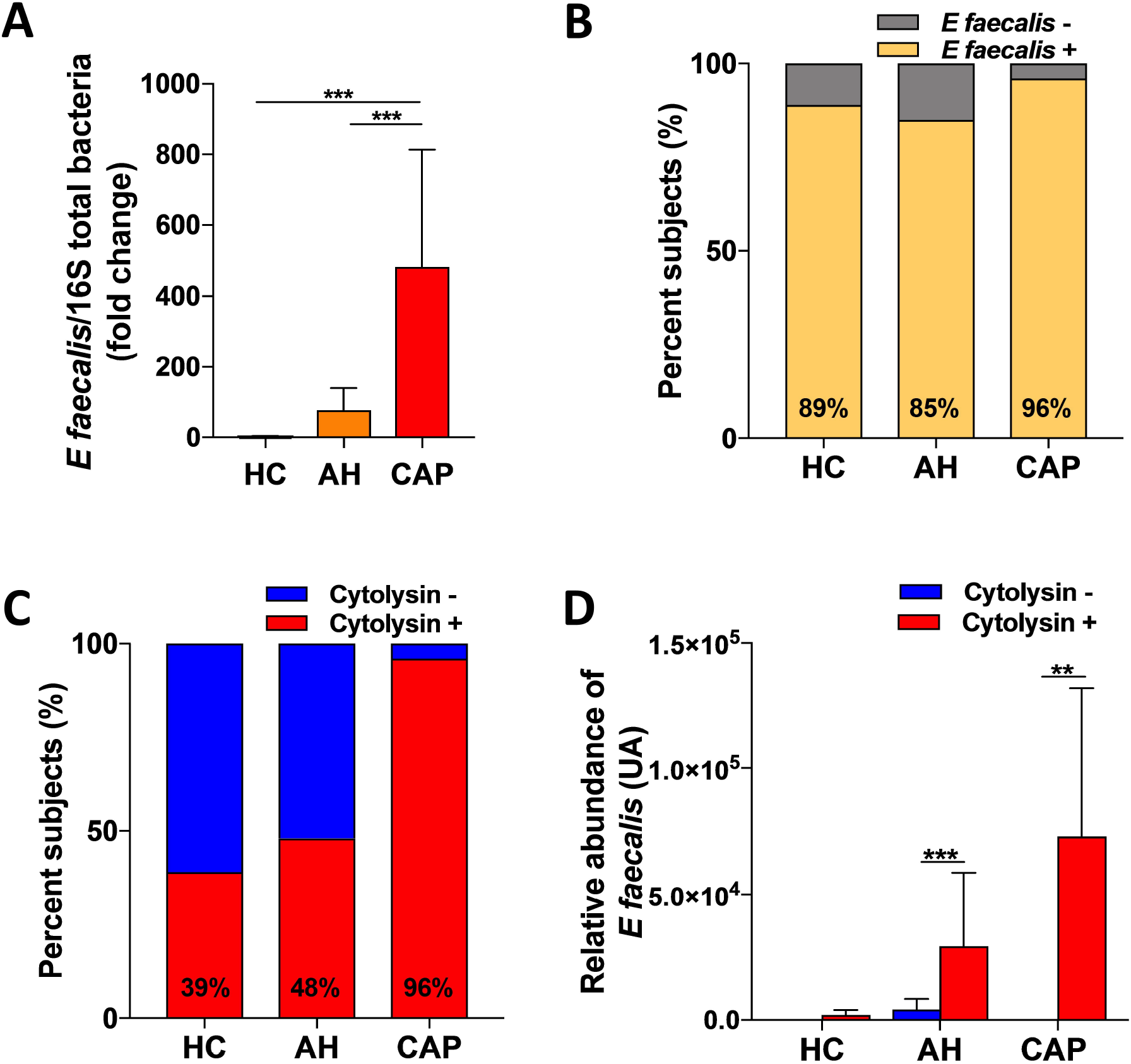
Fecal samples from CAP patients contain more *E. faecalis* and cytolysin than those of AH patients and HC. CAP: chronic alcoholic pancreatitis, AH: severe alcoholic hepatitis, HC: healthy controls. **(a)** Fold change of *E. faecalis* relative abundance in fecal samples from HC (n = 28) and AH (n = 27), and CAP patients (n = 24), assessed by qPCR. **(b)** Percentage of fecal samples positive for *E. faecalis* in HC (n = 28) and AH (n = 27) and CAP patients (n = 24), assessed by qPCR. **(c)** Percentage of cytolysin positive fecal samples in HC (n = 28) and AH (n = 27) and CAP patients (n=24), assessed by qPCR. **(d)** Relative abundance of *E. faecalis* in fecal samples from AH and CAP patients and HC whose fecal samples were cytolysin-positive or cytolysin-negative, assessed by qPCR. Data are shown as the mean ± SEM. Significant results for *p < 0.05, **p < 0.01, and ***p < 0.001 were determined by Mann-Whitney tests, unless stated otherwise.

## DISCUSSION

Overall, our data show that the relative abundance of cytolysin-positive *E. faecalis* is elevated in two different alcohol-related complications i.e. AH and CAP. Therefore, an increase in the relative abundance of cytolysin-positive *E. faecalis* is due to alcohol-consumption itself rather than to a specific complication. Toxicity of cytolytic *E. faecalis* against a human intestinal epithelial cell line has been shown (5) suggesting that it acts directly against the intestinal barrier. Moreover, studies in mice have shown that gut dysbiosis is involved in the pathogenesis of pancreatitis by inducing alterations in intestinal barrier function (6). Cytolysin levels increased significantly in the livers of mice given cytolytic *E. faecalis*, suggesting a direct toxic effect of cytolysin in the liver (3). Therefore, as alcohol itself increases gut permeability, the alcohol-related increase in cytolysin-positive *E. faecalis* in CAP may also be involved in the inflammatory process that leads to CAP. Accordingly, *Enterococcus* is the most common bacteria found in the bile of patients with CAP and antibodies against *E. faecalis* capsular polysaccharide are elevated in the blood of these patients (7). We therefore suggest that targeting cytolysin-positive *E. faecalis* using the bacteriophages described by Duan et al. could also be of great interest for CAP patients.

In summary, we confirm, in our independent French cohort, that patients with AH and no pancreatic complications have an elevated prevalence of cytolysin-positive *E. faecalis* and show that the prevalence of cytolysin-positive *E. faecalis* is also elevated in another alcohol-related complication, CAP without alcoholic AH. This could be of great interest in battling this life-threatening complication, which has no specific treatment and which could benefit from the microbiota-editing strategy using targeted bacteriophages described by Duan et al (3). Further studies are needed to confirm the role of cytolysin-positive *E. faecalis* in CAP and consider it as a therapeutic target in patients.

## Guarantors of the article

Dragos Ciocan, MD, PhD and Anne-Marie Cassard, PhD.

## Specific author contributions

CP: experimental design, acquisition, analysis, and interpretation of data, and drafting of the manuscript. VR, CSV, DC, GP: patients’ recruitment. NT: technical support. GP, VR, AMC, DC: funding. AMC: data analysis and interpretation, drafting of the manuscript. DC: study concept, data analysis and interpretation, drafting of the manuscript. All authors have reviewed and approved the final draft of the manuscript.

## Acknowledgments

The authors thank the Plaimmo Platform, P. Serror (MICALIS, UMT1319 INRA-AgroParisTech) for the generous gift of the *E. faecalis* DNA strain and C Hugot for technical support.

## Financial support

This work was supported by INSERM, Université Paris-Sud, the “Fondation pour la Recherche Médicale” (FRM), the National French Society of Gastroenterology (SNFGE), the “Association Française pour l’Etude du Foie” (AFEF), and the “Groupement Transversal INSERM sur le Microbiote” (GPT microbiota).

## Potential competing interests

The authors declare to have no competing financial interests for the present work.

## Notes

### Competing Interest Statement

The authors have declared no competing interest.

