## SUPPLEMENTARY METHODS for "Cytolysin-positive *Enterococcus faecalis* is elevated in patients with chronic alcoholic pancreatitis"

***Patients.*** Patients with AH were admitted to the Hepato-gastroenterology department of Antoine-Béclère University Hospital, Clamart, France and patients with CAP were recruited at Beaujon University Hospital, Clichy, France. The inclusion and exclusion criteria of patients and healthy individuals has been previously described^2^. General demographic and clinical characteristics were recorded for all patients.

***Bacterial DNA extraction and real-time quantitative qPCR.*** Real-time qPCR was performed using a Light Cycler 480 (Roche Diagnostics) with the LC FastStart DNA Master SYBR Green I kit (Roche Diagnostics). Amplification was initiated with an enzyme activation step at 95°C for 10 min, followed by 40 cycles consisting of a 20-s denaturation step at 95°C, a 15-s annealing step at the temperature appropriate for each primer, and a 45-s elongation step at 72°C. Data were analyzed using Light Cycler 480 Software (Roche Diagnostics). Amplicon quality was verified by gel electrophoresis. Relative gene expression of *E. faecalis* was normalized to that of the 16S bacterial gene.

***Primer sequences.*** The following primer sequences were used: **16S total bacteria forward** (F) 5’-GTGSTGCAYGGYTGTCGTCA-3’, reverse (R): 5’-ACGTCRTCCMCACCTTCCTC-3’ (S=C or G; Y=C or T; M=A or C); ***E. faecalis*** F: 5’-CGCTTCTTTCCTCCCGAGT-3’, R: 5’-GCCATGCGGCATAAACTG-3’, and ***CylL_L_*** F: 5’-CTGTTGCGGCGACAGCT-3’, R: 5’-CCACCAACCCAGCCACAA-3’.

***Statistical analyses.*** Results are presented as the mean ± SEM. Statistical comparisons were performed using unpaired Mann-Whitney, unpaired t-tests, Kruskal-Wallis, ANOVA, chi2 or Fisher exact tests as appropriate (Graphpad Prism, Graphpad Software Inc, La Jolla, California, USA); p < 0.05 was considered to be statistically significant. ^*^p < 0.05, ^**^p < 0.01, ^***^p < 0.001.
